# Functional channels in mature *E. coli* colonies

**DOI:** 10.1101/851428

**Authors:** Liam M. Rooney, William B. Amos, Paul A. Hoskisson, Gail McConnell

## Abstract

Biofilms are important in medicine, industry and the natural environment, however their structure is largely unexplored across multiple spatial scales. We have studied the architecture of mature *Escherichia coli* macro-colony biofilms by means of the Mesolens, an optical system which uniquely allows simultaneous imaging of individual bacteria and over one hundred cubic millimetres of its biofilm milieu. Our chief finding is the presence of intra-colony channels on the order of 10 μm in diameter in *E. coli* biofilms. These channels have a characteristic structure and reform after total mechanical disaggregation of the colony. We present evidence that the channels transport particles and function to assist the absorption of nutrients. These channels potentially offer a new route for the delivery of dispersal agents or antimicrobial drugs to biofilms, ultimately lowering their impact on public health and industry.

## Introduction

Though often inconspicuous, biofilms are one of the most prolific and metabolically active environments on Earth. Biofilms are aggregate communities of microbes held together by extracellular matrices containing extracellular polysaccharides (EPS) and nucleic acids^1,2^. These microbial communities can be composed of one or more species (mono/poly-microbial) and are found in almost every ecological niche^3^. The protective matrix enveloping the biofilm confers resistance to desiccation and exposure to diffusing agents such as biocides or antibiotics^4–8^, in turn promoting the development and spread of antimicrobial resistance^9^. Consequently, the study of biofilm structure is vital to understanding and combatting the development of resistance and lowering the clinical and industrial burden of biofilms. The 3D organisation of biofilms can take many forms^10–13^; for example, mushroom-shaped biofilms grown in liquid flow systems, thin sheet-like biofilms in static liquid systems, pellicle biofilms grown at liquid/air interfaces, and macrocolony biofilms grown on solid surfaces. Although morphologically distinct, what classifies these structurally-different communities as ‘biofilms’ lies with their shared fundamental biochemical signals and pathways^3^.

Dynamic computational modelling programmes, such as CellModeller^14,15^, have been routinely used to predict the spatial patterning and arrangement of cells within bacterial communities^14–18^. *In silico* models primarily show growth of polymicrobial communities where cell shape, size, surface properties and cell-cell interactions influence the spatial organisation of the mature biofilm, resulting in sectoring of different strains into distinct populations, which has been validated experimentally^19–23^. However, *in silico* modelling has shown little evidence of structural ordering or complex spatial patterning, and a lack of effective multi-scale imaging techniques has presented little experimental evidence of 3D structure within mono-strain biofilms.

There has been much imaging of living biofilms. For example, the density-dependent phage sensitivity in *Escherichia coli* colonies has been studied^24^, the biofilms present on human tooth enamel have been imaged at different pH levels^25^, and synchronies of growth and electrical signalling between adjacent bacterial colonies have been observed^26^. Such studies have exposed a gap in the repertoire of the optical microscope in that either microbes could be individually imaged with a high-power objective lens, or the overall structure could be viewed at low magnification with resolution so poor, particularly in depth, that individual microbial cells could not be seen. To address this, we use the Mesolens to image intact live macro-colony biofilms *in situ* with isotropic sub-cellular resolution. In essence, the Mesolens is a giant objective lens with the unique combination of 4× magnification with a numerical aperture (NA) of 0.47; which is approximately five-times greater than that of a conventional 4× objective lens^27^. The low magnification coupled with a high NA result in a field of view (FOV) measuring approximately 6 mm^2^ with lateral resolution of 700 nm and 7 μm axially, while the lens prescription provides a working distance of 3 mm. Moreover, the lens is chromatically corrected across the visible spectrum and designed to be compatible with various immersion routines. While the Mesolens has proven to be a powerful tool in neuroscience, developmental biology and pathology^27–29^, it also presents an untapped technology for biofilm imaging, where we can image whole live microbial communities with unprecedented detail within a single dataset without additional processing or stitching and tiling.

We explored the internal architecture of mature *E. coli* macro-colony biofilms using a novel mesoscopic imaging approach. We identified and characterised a previously undocumented channel system within *E. coli* biofilms. These channels allow for nutrient uptake from the external environment, which offers a novel mechanism for nutrient delivery in microbial communities beyond passive diffusion; which is widely accepted as the main route of delivery for any external compounds to enter a biofilm, whether they be nutrients or antimicrobial drugs^15,16,14,17^. Using fluorescent probes, we determined that intra-colony channels have a significant protein component. Additionally, we demonstrated that intra-colony channels form as an emergent property of biofilm formation in *E. coli*. These findings provide novel understanding of how spatial organisation in bacterial biofilms contributes to their ability to transport material from the external environment – a function which could be exploited to target biofilm dispersal agents and antimicrobial drugs to lower the burden of biofilms on public health.

## Results

### Identification of a network of intra-colony channels in *E. coli* biofilms

We investigated the internal architecture of *E. coli* macro-colony biofilms using conventional widefield epi-fluorescence microscopy, widefield mesoscopy and confocal laser scanning mesoscopy. Using widefield mesoscopy we discovered that *E. coli* (JM105) biofilms contain a network of channel-like structures which permeate the biofilm and travelled from the centre to the leading edge. The channels measure approximately 15 μm wide and appear as non-fluorescing regions within the biofilm which are lined by individual cells in a pole-to-pole arrangement. We applied a Classic Maximum Likelihood Estimation deconvolution algorithm to a z-stack acquired using the Mesolens in widefield epi-fluorescence mode to improve image quality and reveal the arrangement of individual cells in a mature macro-colony biofilm. We then applied a colour-coded look-up table (LUT) according to the axial position of each optical section within the 36 μm-thick z-stack (Figure 1). From the axial-coded LUT we can see that the intra-colony channels are not merely 2D lateral arrangements of cells, but that the channels have a 3D topography within the context of the biofilm, resembling canyons and ravines rather than enclosed capillaries.

**Figure 1.**
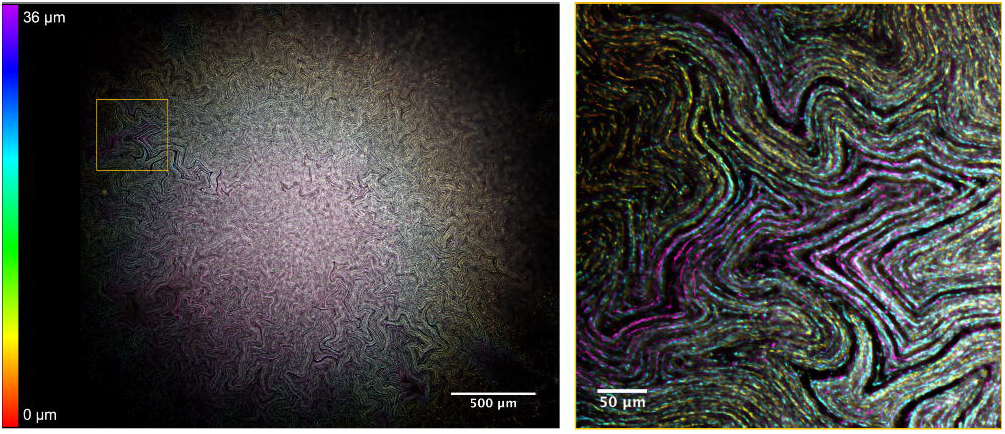
Visualising the intra-colony channel system of *E. coli* macro-colony biofilms. A deconvolved 36 μm-thick transverse sub-stack of a mature *E. coli* macro-colony biofilm acquired using widefield mesoscopy. An axial colour-coded LUT has been applied which indicates the relative position of each cell within the context of the biofilm. A magnified ROI is presented where individual cells can be clearly resolved. Channel structures are seen to permeate throughout the biofilm and present a 3D topography within the context of the biofilm.

Additionally, we imaged JM105 biofilms using confocal mesoscopy to ensure that the deconvolution algorithm that we used when processing widefield Mesolens data did not introduce erroneous structural artefacts. Confocal microscopy provides a marked improvement in signal-to-noise ratio compared to widefield techniques, particularly with thick specimens, resulting in a similar image quality to a deconvolved widefield dataset. Confocal mesoscopy revealed the same channel structures that we identified in widefield imaging experiments presented in Figure 1 (Supplementary Figure 2, Supplementary Movie 1). This concludes that the structures we observed were not introduced as an artefact of image processing.

To demonstrate the benefit of using the Mesolens over conventional microscopes for imaging live biofilms, we also imaged biofilms using a conventional upright widefield epi-fluorescence microscope with a low-magnification, low-NA lens (4×/0.13 NA). We compared the ability of the Mesolens and the conventional microscope to resolve the intra-colony channels and found that there was a clear improvement in the spatial resolution with the Mesolens (Supplementary Figure 3). The resolution improvement applies to both lateral and axial resolution, and establishes the Mesolens as an ideal imaging technology for 3D imaging of large microbiological specimens with sub-micron resolution.

### A structural assessment of intra-colony channels

The channel structures we have identified appear as dark regions within the biofilm, and so we hypothesised that they contained some form of structural matrix. We began investigating the structural makeup of the channels by determining if they were filled with materials of differing refractive index compared to that of the biomass. Possible candidates were solid growth medium which was forced upward during biofilm growth, or air which was trapped in the biofilm. We used reflection confocal mesoscopy where signal is detected from reflections of incident light at refractive index boundaries, such as those between bacterial cells and the surrounding growth medium. A maximum intensity projection of an unlabelled *E. coli* JM105 biofilm acquired in reflection confocal mode showed no reflection signal resembling the intra-colony channels which we report (Figure 2a). This informs us that the channels must be of a similar refractive index to the surrounding biomass and biofilm matrix and are not occupied by solid growth medium or air.

To determine if the channel structures we observe were occupied by non-viable/non-fluorescing cells, biofilms were grown in the presence of the viability dye, Sytox Green. This dye has an emission peak at 523 nm enabling the use of HcRed1 (*λ*_*em*_. 618 nm) expressing JM105 *E. coli* cells for two-colour imaging. Figure 2b shows a false-coloured composite maximum intensity projection of a JM105-miniTn7-*HcRed1* biofilm stained with Sytox Green acquired using widefield mesoscopy, where live cells are presented in cyan and non-viable cells are shown in yellow. We subtracted the signal of the non-viable cells from HcRed1-expressing cells to prevent spectral overlap in the emission of the two fluorophores, meaning that no Sytox-labelled cells were falsely presented in cyan. We observed that viable and non-viable cells formed two distinct domains within the colony. Here, non-viable cells cluster in the centre of the biofilm while intra-colony channels are not occupied by non-viable/non-fluorescent cells.

**Figure 2.**
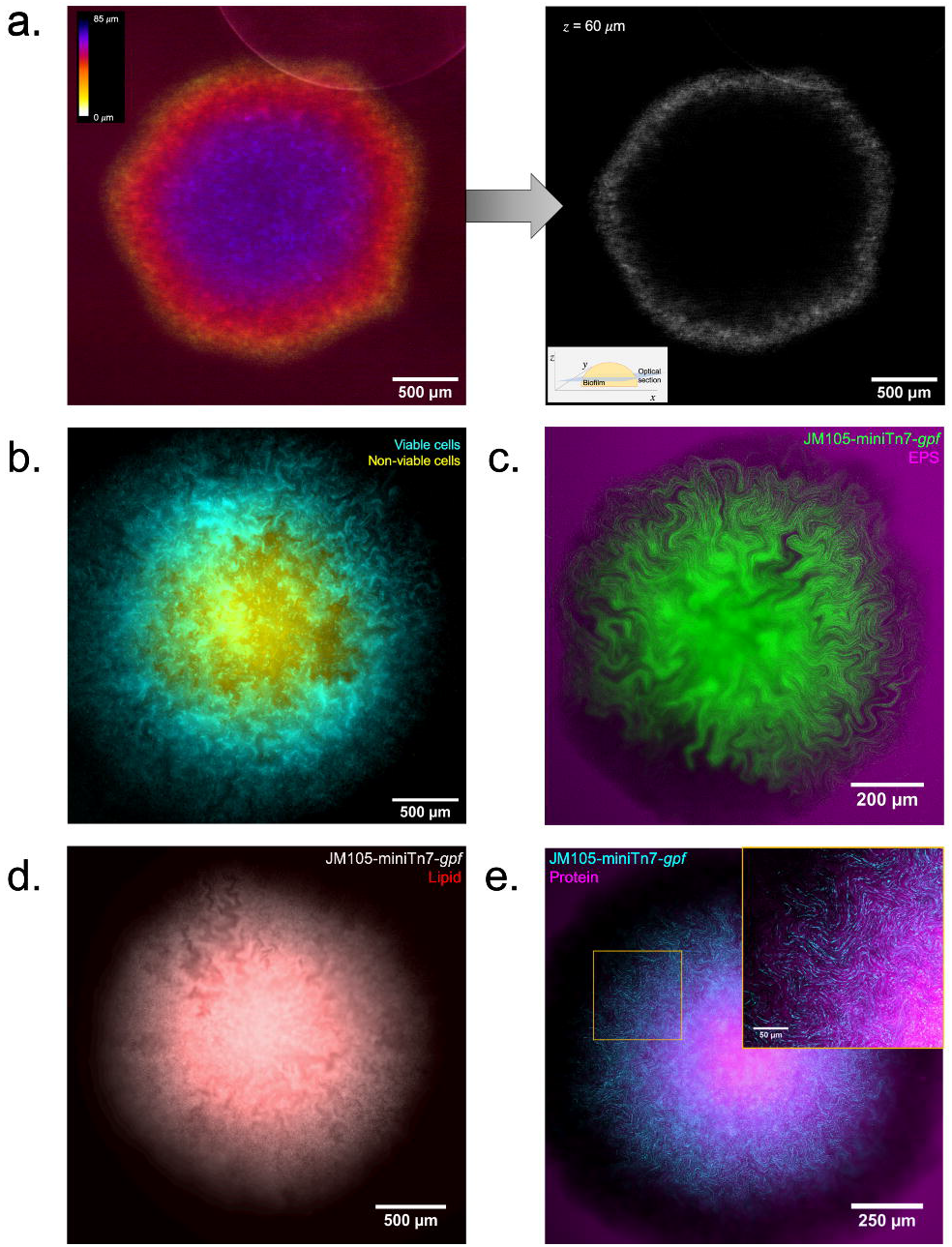
Characterising the structure of intra-colony channels. **(a)** Maximum intensity projection of an unlabelled JM105 colony acquired using reflection confocal mesoscopy, with a single isolated optical section shown. Reflection imaging determined that intra-colony channels were not occupied by material of differing refractive index to the biomass. The colony-medium interface can be observed clearly, while there is no evident structure within the colony. **(b)**. Signal from non-viable cells (yellow) was subtracted from viable cells to negate any spectral overlap in the emission of Sytox Green and HcRed1. A composite maximum intensity projection of the entire colony is presented. Intra-colony channels in the viable cell population (cyan) did not contain any non-viable cells. **(c)** Alexa594-WGA-stained EPS residues (magenta) were not present in the intra-colony channels when compared with elsewhere in the biofilm, meaning channels were not composed of an EPS-based matrix. **(d)** Nile Red-stained lipids (red) clustered in the centre of *E. coli* biofilms while intra-colony channels remain unstained by Nile Red. Therefore, intra-colony channels were not composed of lipids. **(e)** Emission of SYPRO Ruby-stained extracellular proteins (magenta) mimicked the spatial patterns of intra-colony channels, showing that channels were filled by a protein-based matrix.

To investigate whether intra-colony channels were filled with exopolysaccharides (EPS) secreted by bacteria within the biofilm, we grew JM105 biofilms in the presence of the lectin binding dye conjugate Alexa594-Wheat germ agglutinin (WGA). Figure 2c shows a deconvolved composite image of a JM105-miniTn7-*gfp* biofilm (green) and associated EPS (magenta). Exopolysaccharides are distributed throughout the entire biofilm and are not strictly localised within the channel structures. We assessed lipid distribution using the lipid-binding dye Nile Red, which showed that intra-colony channels are not composed of a lipid matrix (Figure 2d). Extracellular proteins were stained using the protein-specific fluorescent dye, SYPRO Ruby. The intra-colony channels contained extracellular protein (Figure 2e). The finding that the channels are filled by a protein matrix suggests that they arise not due to some stochastic process and that the channels have some function.

### Channels emerge as an inherent property of biofilm formation

To determine whether the formation of intra-colony channels arose as an emergent property of biofilm formation, we investigated if the structures were able to re-form following disruption. We allowed the biofilm to establish the formation of channels (Figure 3a) and then disturbed the colony, by mixing to create a uniform mass of cells. Following a recovery period of 10 hours, the channels reformed in the regrowing regions of the biofilm (Figure 3b). The ability of the channels to form in the same way as a naïve colony suggests that they form as an emergent property of *E. coli* colonial growth on a solid surface.

**Figure 3.**
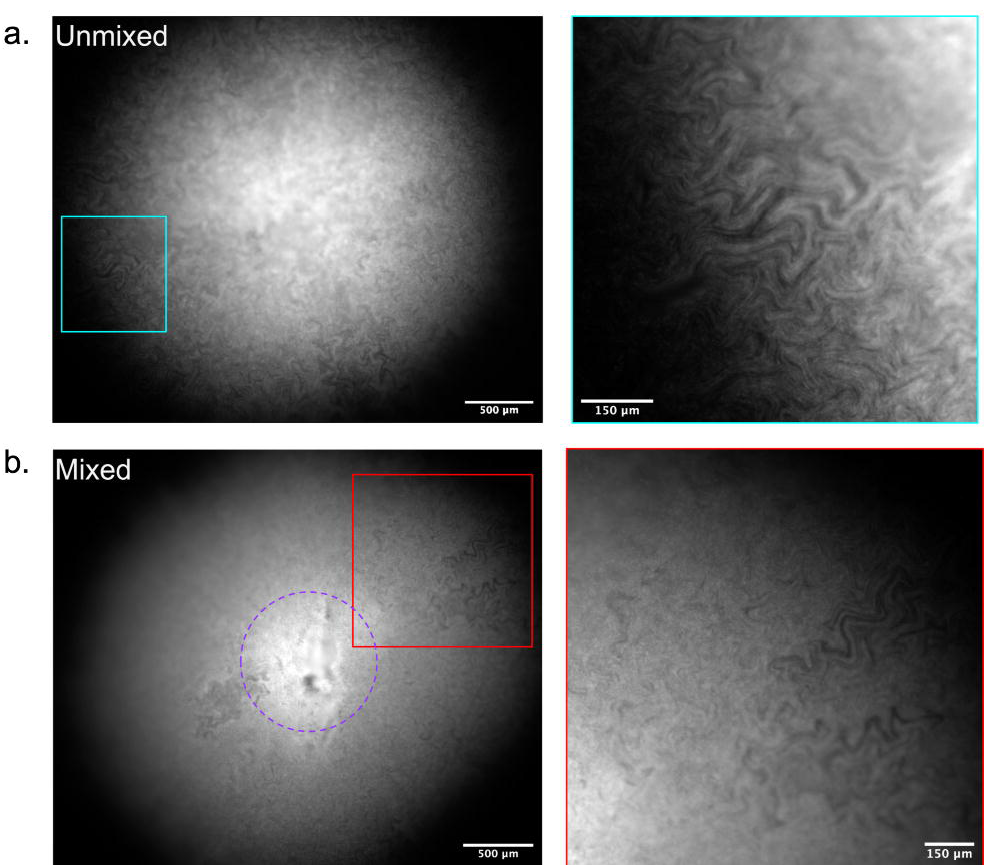
Intra-colony channels form as an emergent property of biofilm formation. **(a)** An unmixed, naïve control biofilm of JM105-miniTn7-*gfp* with established intra-colony channels. **(b)** A macro-colony JM105-miniTn7-*gfp* biofilm which was initially grown for 10 hours before mechanical disruption and subsequent recovery and regrowth at 37°C for a further 10 hours. Regrowth was accompanied with the re-emergence of intra-colony channels in the outgrown region of the disrupted colony, showing that channel formation is an emergent property of macro-colony biofilm development.

### Channels are unable to cross strain boundaries in mixed isogenic cultures

Growth of two isogenic strains in co-culture, each expressing a different photoprotein, two strains formed sectors as has been previously described ^18–20,22,21,23^. We wished to explore this sectoring property in the context of intra-colony channel formation and to determine if the channels were shared between the strains. When the two isogenic strains sector, the channels do not intersect the boundary between the strains and are retained within the own sector (Figure 4). The confinement of channels was more evident between different populations (i.e. HcRed1 and GFP-expressing), whereas the boundaries between sectors of cells expressing the same photoprotein were less ordered.

**Figure 4.**
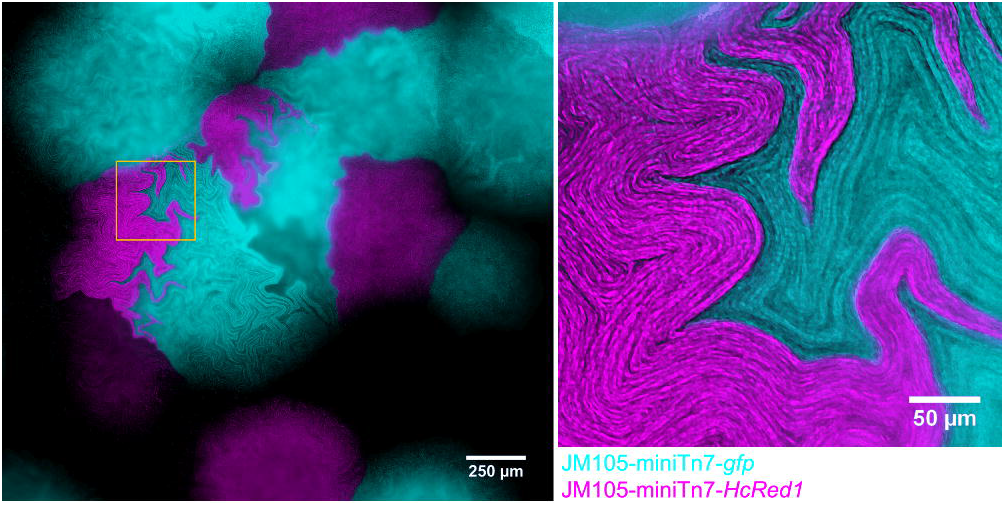
Intra-colony channels are confined within clonal populations and unable to cross strain boundaries. A mixed culture of isogenic JM105 strains which express either GFP (cyan) or HcRed1 (magenta). Each strain sectored into segregated clonal populations which have propagated from a single colony forming unit, and cells from each sector were unable to cross the strain boundary. The intra-colony channels present within each sector were also unable to cross the strain boundary and were therefore not shared by opposing isogenic colonies.

### Intra-colony channels present a novel nutrient acquisition system in *E. coli* biofilms

To investigate whether the intra-colony channels play a role in the transport of substances into the biofilm, the functional role of the channel system was tested by introducing 200 nm diameter fluorescent microspheres to the extracellular medium when preparing the specimen for widefield mesoscopy. The fluorescent microspheres were spread as a dense lawn along with a dilute mid-log JM105-miniTn7-*gfp* culture. A single optical section, 25 μm above the base of the colony, allows the outline of the colony to be observed at the edges of the image, with the untouched lawn of microspheres outside the colony (Figure 5a). The distribution of beads in these areas are homogenous, whereas within the colony the transport of the fluorescent microspheres through the channels reflects the spatial structure of the biofilm. Magnified regions of interest (ROI) of intra-colony channels show that the channels are acting as conduits for the transport of microspheres into the biofilm (Figure 5b). The transport of microspheres into the channels suggests that these intra-colony structures are involved in the acquisition of substances from the external environment. We suggest the ability of channels to transport small fluorescent particles could be extended to facilitate uptake of smaller particles into the colony, such as nutrients.

**Figure 5.**
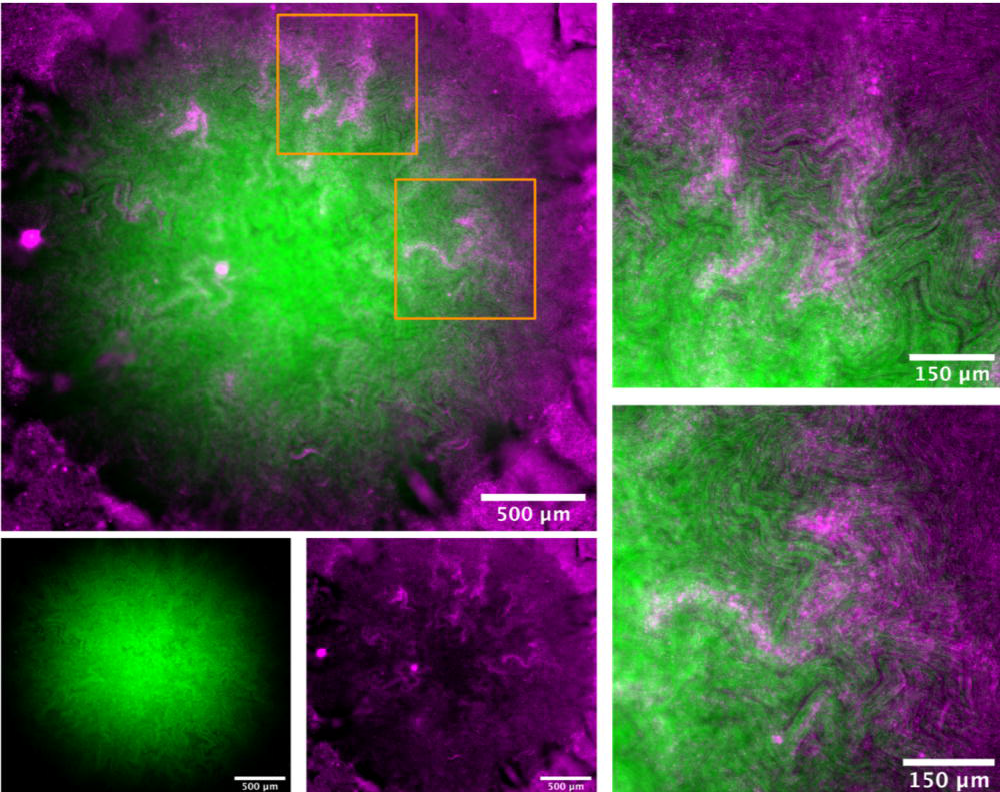
Intra-colony channels facilitate transport of microscopic particles. A single optical section approximately 25 μm above the base of the colony shows a mature JM105-miniTn7-*gfp* biofilm (green) and a lawn of 200 nm fluorescent microspheres (magenta). The fluorescent microspheres were transported from a confluent lawn at the base of the colony into the intra-colony channels and directed towards the centre of the colony. Two ROIs are presented from different regions of the colony where fluorescent microspheres were transported into the colony via intra-colony channels.

To further investigate the role of intra-colony channels in biofilm nutrient acquisition, an arabinose inducible GFP strain (*E. coli* JM105 P_BAD_*-gfp*) was used. Growth of the arabinose-inducible GFP strain on solid minimal medium with L-arabinose as the sole carbon source revealed the biofilm fluoresced most intensely in regions which bordered the intra-colony channels (Figure 6). This suggests that the concentration of L-arabinose is highest in the channel system compared to the remainder of the biofilm and demonstrates the role of these structures in nutrient acquisition and transport within the colony. This finding challenges the long-held belief that bacterial colony nutrient uptake occurs through simple diffusion through the extracellular matrix of the biofilm, and concurs with previous work which suggested that large biofilms must develop transport mechanisms to direct nutrients to their centre^1^.

**Figure 6.**
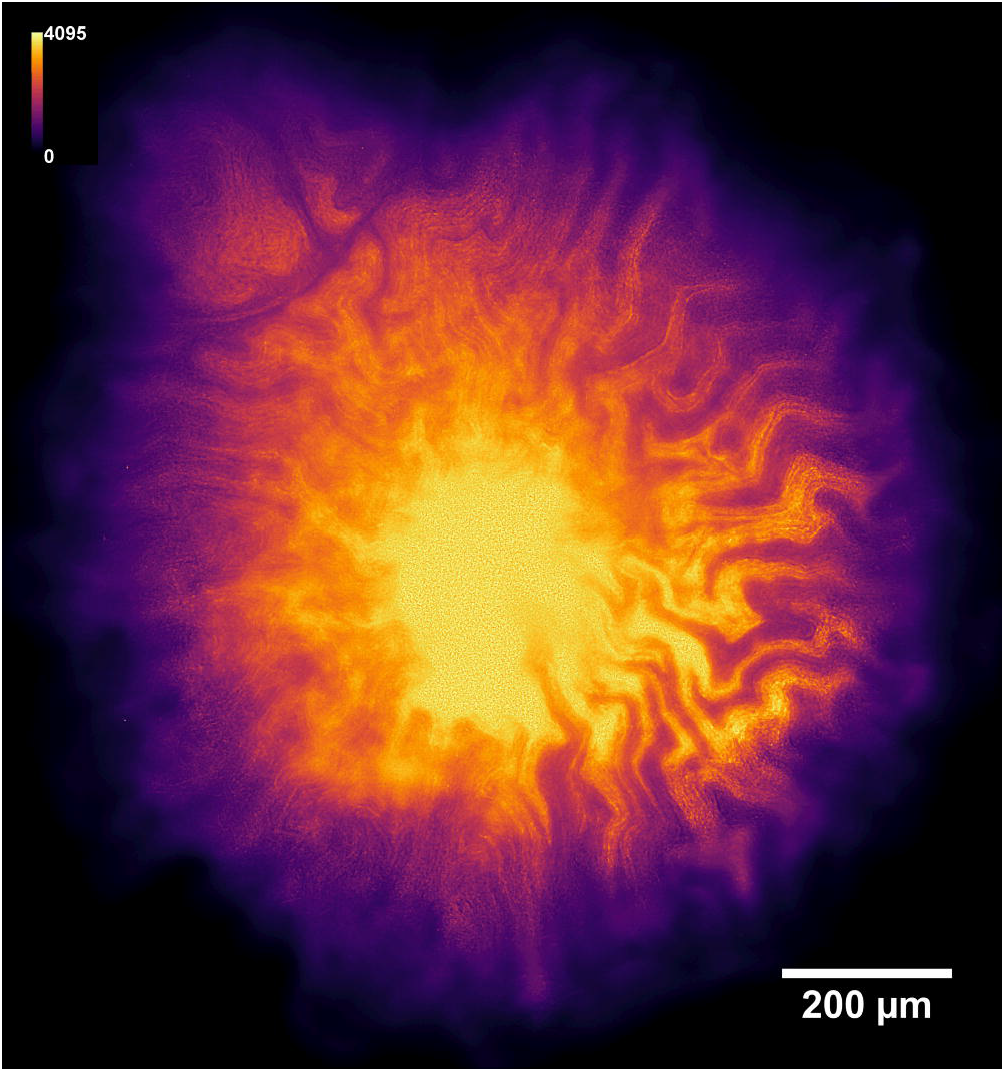
Intra-colony channels play a functional role in nutrient acquisition and transport to the centre of bacterial biofilms. A deconvolved image of a JM105-pJM058 macro-colony biofilm grown on M9 minimal medium with L-arabinose as the sole carbon source. This arabinose biosensor expresses GFP only in the presence of L-arabinose. GFP emission intensity was higher in cells which line the intra-colony channels compared to cells elsewhere within the biofilm, which shows that the channel structures have a higher concentration of L-arabinose compared to elsewhere within the biofilm. This provides evidence of a functional role in nutrient acquisition and transport for the intracolony channel system.

## Discussion

This study is the first application of the Mesolens to microbiology and has offered a new approach for imaging large microbial specimens, enabling us to characterise a novel structural aspect of *E. coli* macro-colony biofilms. The channel structures reported are formed as an emergent property of biofilm growth and are confined within founder cell boundaries in mixed isogenic cultures. We have also established a functional role for intra-colony channels in nutrient acquisition and transport. The identification and characterisation of these previously undocumented channels offers a novel outlook for microbial community biology and provides a novel mechanism for nutrient delivery in bacterial biofilms.

Previous biofilm imaging studies have mainly used conventional widefield and laser scanning microscopy to study biofilm architecture, which are inherently limited by sacrificing spatial resolution and imaging volume. For example, automated tile-scanning microscopes which change the location of the FOV or focal plane have been used to image growing colonies from 1×10^1^−1×10^4^ cells^30–32^; however, this method often requires long acquisition periods and results in tiling artefacts. With the Mesolens we negate the need for stitching and tiling when imaging multi-millimetre specimens and can image beyond small bacterial aggregates to visualise live bacterial macro-colonies in excess of 1×10^9^ cells while maintaining sub-micron resolution throughout the entire 6 mm^2^ field. Therefore, in comparison with other conventional large specimen imaging techniques, the Mesolens stands as a novel and improved method for *in situ* imaging of live bacterial communities. Additionally, recent advances in light sheet microscopy^33^ and mutli-photon microscopy^34,35^ have been applied to biofilm imaging. However, these methods currently cannot resolve sub-micron information over multi-millimetre scales, as with the Mesolens. The same problem accompanies ultrasound^36,37^, optical coherence tomography and photoacoustic tomography^38–40^ methods used for mesoscale biofilm imaging, where they cannot properly resolve structures on the order of which we report. We have also studied images of bacterial macro-colonies under a widely available conventional stereomicroscope. Careful comparison with Mesolens images suggests that traces of the channel may be faintly visible in spite of the low resolution of stereomicroscopes in x, y and particularly z-dimensions.

The structures we have identified bear similarities to some other aspects of bacterial community architecture, however it is important to note that the channels we identify are fundamentally different to structures such as the water irrigation channels discovered in mushroom-shaped *Pseudomonas* and *Klebsiella spp*. biofilms^41,42^. There have also been channel-like structures identified in mature bacterial colonies, such as the crenulations of *B. subtilis* macro-colonies^43,44^ or the macroscopic folds of *P. aeruginosa* biofilms^45,46^, which have been extensively described in literature. It is important to note, that crenulations and folds are all visible as surface structures of the colony and resolvable using photography techniques, whereas the intra-colony channels identified here are present within the main body of the biofilm and are not observable by viewing the surface of the colony. A similar phenomenon was recently reported in colonies of *Proteus mirabilis* where 100 nm diameter fluorescent microspheres were observed to penetrate the boundary of the colony through “crack-like conduits” present at the colony edge^55^. However, the authors were unable to show any spatial evidence of the conduits themselves.

The spatial arrangement of the intra-colony channels is fractal in nature, with repeating patterns and complex topographies. Upon first glance, channels resemble fractal features found in multi-strain colonies which form as a result of the mechanical instability between growth and viscous drag of dividing cells^19^. However, these features have only been reported in multi-strain colonies where the fractal dendrites have been composed of live, fluorescing cells^20–23,47^. We demonstrate that the spatial patterns we observe are different to those outlined previously. Firstly, the patterns we observe arise in a single population of cells where there are no strain-to-strain interactions to result in the formation of fractal patterns. Given that the intra-colony channels are not occupied by dead non-fluorescing cells (Figure 2b) it is clear that the bacterial colonies used in this work are not composed of two pseudo-domains (i.e. viable and non-viable cells) which could interact to form complex 3D fractal patterning. Our finding that non-viable cells localise in the centre of the biofilm agrees with previous studies showing that dense microbial aggregates often have dense hypoxic, acidic centres which have diminished access to nutrients^11,12,34,48–52^.

The intra-colony channels form as an inherent property of biofilm formation, leading to fractal-like patterns which exhibit plasticity which is reminiscent of the results of a classical eukaryotic developmental biology experiment by Moscona, where reformation of the channel architecture in marine sponges occurred after disaggregation by passage through a fine silk mesh^53,54^. The ability of the channels to reform also suggests that they fulfil a functional role in the context of the biofilm.

In summary, we have identified a previously undocumented nutrient uptake system in colonial biofilms which challenge the current belief that cells which are out with the reach of underlying nutrient-rich medium are able to gain nutrients beyond simple diffusion through the base of the biofilm^14–17^. The presence of these channels may represent a route to circumvent the chemical protection and resistance phenotype of bacterial biofilms^56^, such that rather than applying drugs to the apical surface of the biofilm it may be possible to exploit the intra-colony channels for delivery of antimicrobial agents. The identification and characterisation of an intra-colony channel network could therefore have far-reaching applications to public health and disease prevention, while providing another understanding on the delivery of nutrients to the centre of densely packed microbial communities.

## Materials and Methods

### Designing and 3D-printing a chamber slide for biofilm imaging

A custom imaging chamber was designed using AutoCAD (Autodesk, USA) with the purpose of imaging large-scale cultured bacterial communities *in situ* using the Mesolens. The design consisted of a plate with dimensions 90 mm × 80 mm × 12 mm and a central well measuring 60 mm in diameter with a depth of 10 mm (Supplementary Figure 1). The imaging chamber was 3D-printed using black ABS plastic (FlashForge, Hong Kong) with a FlashForge Dreamer 3D printer (FlashForge, Hong Kong). The chamber slide was sterilised prior to use with 70% ethanol and UV irradiation for 15 minutes.

### Bacterial strains and growth conditions

All experiments were performed using the *E. coli* strains outlined in Supplementary Table 1. Colony biofilms were grown by inoculating a lawn of cells at a density of 1×10^4^ cfu/ml on either solid LB medium or M9 minimal medium^57^ supplemented with the appropriate selective antibiotic to achieve single colonies. The colonies were grown in the 3D-printed imaging mould at 37°C for 18-24 hours in darkened conditions prior to imaging.

### Specimen preparation

For colony imaging alone, colonies were submerged in sterile LB broth (refractive index (*n*) = 1.338) as a mounting medium following the allocated growth time prior to imaging. A large coverglass was placed over the central well of the imaging mould (70 mm × 70 mm, Type 1.5, 0107999098 (Marienfeld, Lauda-Koenigshofen, Germany)), and the colonies were then imaged using either the Mesolens or a conventional widefield epi-fluorescence microscope to compare their performance and to justify using the Mesolens to study biofilm architecture over conventional techniques.

The refractive index of the LB mounting medium was measured using an Abbe Refractometer (Billingham & Stanley Ltd., U.K.) which was calibrated using Methanol at 21°C.

### Conventional widefield epi-fluorescence microscopy

Colony biofilms were imaged on a conventional an Eclipse E600 upright widefield epi-fluorescence microscope (Nikon, Japan) equipped with a 4×/0.13 NA PLAN FLUOR objective lens (Nikon, Japan). GFP excitation was provided by a 490 nm LED from a pE-2 illuminator (CoolLED, U.K.), and emission was detected using a bandpass filter (BA 515-555 nm, Nikon, Japan) placed before an ORCA-spark digital CMOS camera (Hamamatsu, Japan). The camera detector was controlled using WinFluor software^58^. Colonies were imaged after 20 hours of growth in an imaging mould as described above.

### Widefield epi-fluorescence mesoscopy

Specifications of the Mesolens have been previously reported^27^, and therefore only the imaging conditions used in this study will be outlined here. GFP excitation was achieved using a 490 nm LED from a pE-4000 LED illuminator (CoolLED, U.K.). A triple bandpass filter which transmitted light at 470 ± 10 nm, 540 ± 10 nm and 645 ± 50 nm was placed in the detection pathway. The emission signal was detected using a VNP-29MC CCD camera with chip-shifting modality (Vieworks, South Korea) to capture the full FOV of the Mesolens at high resolution. Widefield mesoscopic imaging was carried out using water immersion (*n* = 1.33) with the Mesolens’ correction collars set accordingly to minimise spherical aberration through refractive index mismatch.

### Confocal laser scanning mesoscopy

For laser scanning confocal mesoscopy specimens were prepared as outlined above. Fluorescence excitation of GFP was obtained using the 488 nm line set at 5 mW from a multi-line LightHUB-4 laser combiner (Omicron Laserage, Germany). The green emission signal was detected using a PMT (P30-01, Senstech, U.K.) with a 550 nm dichroic mirror (DMLP550R, Thorlabs, USA) placed in the emission path and a 525/39 nm bandpass filter (MF525-39, Thorlabs, USA) placed before the detector.

For reflection confocal mesoscopy incident light was sourced from a 488 nm line set at 1 mW from a multi-line LightHUB-4 laser combiner (Omicron Laserage, Germany). Reflected signal was detected using a PMT (P30-01, Senstech, U.K.) with no source-blocking filter in place.

Confocal laser scanning mesoscopy was carried out using type DF oil immersion (*n* = 1.51) with the Mesolens’ correction collars set accordingly to minimise spherical aberration through refractive index mismatch.

### Structural assessment of intra-colony channels

To characterise the structure of intra-colony channels we sought to visualise the distribution of several archetypal structural components of biofilms.

As the biofilms in this study were submerged during imaging in a medium with known refractive index, we were able to determine if channels were filled with substances of differing refractive index (e.g. air) using reflection confocal mesoscopy as above. Solid LB was cast into a 3D printed imaging chamber and inoculated with JM105 at a density of 1×10^4^ cfu/ml and incubated for 18-24 hours at 37°C in darkened conditions. Biofilms were mounted in sterile LB medium (*n* = 1.338) prior to imaging.

We then imaged the distribution of non-viable cells in the biofilm based on the approach developed by Asally^43^. Briefly, JM105-miniTn7-*HcRed1* colony biofilms were grown for imaging in 3D-printed imaging moulds as outlined previously. LB medium was supplemented with gentamicin (20 μg/ml) and 0.5 μM Sytox green dead-cell stain (S7020, Invitrogen, USA). Cells were seeded at a density of 1×10^4^ cfu/ml and grown for 18-24 hours prior to imaging on the Mesolens in widefield epi-fluorescence mode as described above. A 490 nm and a 580 nm LED from a pE-4000 LED illuminator (CoolLED, U.K.) were used to excite Sytox Green and HcRed1 respectively. The emission signal was detected using a VNP-29MC CCD detector (Vieworks, South Korea) with 3×3 pixel-shift modality enabled and with a triple band pass filter (470 ± 10 nm, 540 ± 10 nm and 645 ± 50 nm) in the emission path.

To visualise the distribution of EPS in the biofilm we stained sialic acid and *N*-acetylglucosaminyl residues by supplementing solid M9 medium (0.2% glucose (w/v))^57^ with 20 μg/ml gentamicin and 2 μg/ml Alexa594-WGA (W11262, Invitrogen, USA) before inoculating with 1×10^4^ cfu/ml JM105-miniTn7-*gfp* and growing as previously described. We imaged EPS-stained specimens using widefield epi-fluorescence mesoscopy as before using a 490 nm LED to excite GFP and 580 nm LED to excite Alexa594-WGA.

We determined the lipid localisation throughout the biofilm by staining with Nile Red. We supplemented solid LB medium with 20 μg/ml gentamicin and 10 μg/ml Nile Red (72485, Sigma-Aldrich, USA) before inoculating with 1×10^4^ cfu/ml JM105-miniTn7-*gfp* and growing as previously described. We then imaged the lipid distribution in relation to the intra-colony channels using widefield epi-fluorescence mesoscopy as before using a 490 nm LED to excite GFP and 580 nm LED to excite Nile Red.

The protein distribution was determined by staining the biofilm with FilmTracer SYPRO Ruby biofilm matrix stain (F10318, Fisher Scientific, USA) which binds to a number of different classes of extracellular protein. Solid LB medium was prepared containing 20 μg/ml gentamicin and a final concentration of 2% (v/v) FilmTracer SYPRO Ruby biofilm matrix stain before inoculating with JM105-miniTn7-*gfp* and growing as previously described. Specimens were imaged using widefield epi-fluorescence mesoscopy. A 490 nm and a 580 nm LED from a pE-4000 illuminator (CoolLED, UK) were used for GFP and SYPRO Ruby excitation, respectively. Fluorescence emission from GFP and SYPRO Ruby were detected as outlined above. Both channels were acquired sequentially.

### Disruption and recovery of intra-colony channel structures

To assess the ability of the structures we observe to recover following disruption, single colonies of JM105-miniTn7-*gfp* were grown on solid LB medium supplemented with 20 μg/ml gentamicin and allowed to grow for 10 hours at 37°C in darkened conditions. Following the initial growth step colonies were removed from the incubator and gently mixed with a sterile 10 μl pipette tip to disrupt the channel structures in the growing biofilm. Care was taken to prevent disruption to the underlying solid medium on which the colony was supported. Following disaggregation, the colonies were grown for a further 10 hours at 37°C in darkened conditions prior to imaging. Colonies were then mounted in sterile LB medium and imaged using widefield epi-fluorescence mesoscopy as described above.

### Using differentially labelled isogenic strains to observe channels in mixed cultures

The phenomenon of strain sectoring has been previously documented and occurs by mechanical buckling as adjacent colonies expand into each other during radial growth^18,19^. We investigated whether intra-colony channels were able to cross the strain boundary between sectors by inoculating a low-density mixed culture of JM105-miniTn7-*gfp* and JM105-miniTn7-*HcRed1* at a 1:1 ratio and inoculating a lawn onto solid LB medium containing 20 μg/ml gentamicin. We allowed colonies of each strain to stochastically collide into adjacent clonal populations during colony expansion and then imaged using widefield mesoscopy after incubation for 20 hours at 37°C in darkened conditions as described above. We used colony PCR to confirm that the miniTn7 insertion, which contained the photoprotein gene, occurred at the same chromosomal location in both strains (*glmS* Fwd. – 5’ AAC CTG GCA AAT CGG TTA C; *tn7*R109 Rev. – 5’ CAG CAT AAC TGG ACT GAT TTC AG). The miniTn7 transposon inserts at only one *attTn7* site in the chromosome, downstream of *glmS*^59^. We found that both JM105-miniTn7-*gfp* and JM105-miniTn7-*HcRed1* were both inserted approximately 25 base pairs downstream of *glmS*. Therefore, there is no genotypic difference between the strains, save for the inserted photoprotein gene.

### Fluorescent microsphere uptake assay

To assess the function of the structures we observe, a confluent lawn of fluorescent microspheres was seeded along with the bacterial inoculum at the culturing stage. Two-hundred nanometre multi-excitatory microspheres (Polysciences, Inc., USA) were seeded at a density of 1×10^10^ microspheres/ml and plated along with 1×10^4^ cfu/ml JM105-miniTn7-*gfp* in a mixed-inoculum. Microsphere translocation was assessed by widefield epi-fluorescence mesoscopy as above with two-channel detection for both the GFP and microsphere fluorescence emission. A triple bandpass emission filter which transmitted light at 470 ± 10 nm, 540 ± 10 nm and 645 ± 50 nm was place in the detection path. Sequential excitation of GFP and the fluorescent microspheres was achieved using a 490 nm and 580 nm LED, respectively, from a pE-4000 LED illuminator (CoolLED, U.K.) Each channel was acquired sequentially using a CCD camera detector (Stemmer Imaging, U.K.). All imaging was carried out using water immersion.

### Assessing the role of intra-colony channels in nutrient uptake

The functional role of the structures which we observe was tested using an arabinose biosensor where GFP expression was controlled by the presence or absence of L-arabinose. The biosensor strain contained the *araBAD* operon with *gfp* inserted downstream on the promotor and ara*BAD* functional genes. The biosensor strain was a gift from colleagues at the James Hutton Institute.

JM105 transformed with the arabinose biosensor plasmid, pJM058, were grown overnight at 37°C while shaking at 250 rpm in liquid LB medium supplemented with 25 μg/ml chloramphenicol. Overnight cultures were then diluted in fresh LB and grown until OD_600_ = 0.5. Cells were then pelleted and washed three times with 1× M9 salts. Washed cells were inoculated on to solid M9 minimal medium^57^ with L-arabinose as the sole carbon source (0.2%) at a density of 1×10^4^ cfu/ml and grown for 42-48 hours in darkened conditions at 37°C. Specimens were then prepared for imaging as outlined above.

### Image processing and analysis

Widefield epi-fluorescence mesoscopy *z*-stacks were deconvolved where specified using with Huygens Professional version 19.04 (Scientific Volume Imaging, The Netherlands, http://svi.nl) using a Classic Maximum Likelihood Estimation algorithm. A theoretical point spread function was generated using Huygens Professional with parameters adjusted to suit the experimental setup. Deconvolution was performed using a server with a 64-bit Windows Server 2016 Standard operating system (v.1607), two Intel^®^ Xeon^®^ Silver 4114 CPU processors at 2.20 GHz and 2.19 GHz and 1.0 TB installed RAM. Image analysis was performed using FIJI^60^. Figures presented here were linearly contrast adjusted for presentation purposes where required using FIJI^60^.

## Acknowledgements

The authors would like to thank Lee McCann (formerly University of Strathclyde, UK) for his technical input with the Mesolens and help with initiating the experiments. In addition, we would like to thank Ainsley Beaton (University of Strathclyde, UK) for the kind gift of the JM105-miniTn7-*gfp* and JM105-miniTn7-*HcRed1* strains, and to Morgan Feeney (University of Strathclyde) for her advice on this manuscript. We also thank Nicola Holden and Jacqueline Marshall (James Hutton Institute, UK) for the kind gift of the pJM058 plasmid which contained the P_BAD_-*gfp* biosensor. This work was supported by the Medical Research Council (MR/K015583/1).

## Author Contributions

LMR conducted all experiments and analysed all data. LMR, WBA, PAH and GM were responsible for the experimental design. LMR, WBA, PAH and GM prepared the manuscript.

## Materials and Correspondence

Any requests for materials or correspondence should be directed to LMR.

## References

1. Hobley, L., Harkins, C., MacPhee, C. E. & Stanley-Wall, N. R. Giving structure to the biofilm matrix: an overview of individual strategies and emerging common themes. FEMS Microbiol. Rev. 39, 649–669 (2015).

2. Nadell, C. D., Drescher, K. & Foster, K. R. Spatial structure, cooperation and competition in biofilms. Nat. Rev. Microbiol. 14, 589 (2016).

3. Flemming, H.-C. & Wuertz, S. Bacteria and archaea on Earth and their abundance in biofilms. Nat. Rev. Microbiol. 17, 247–260 (2019).

4. Costerton, J. W. et al. Bacterial biofilms in nature and disease. Annu. Rev. Microbiol. 41, 435–64 (1987).

5. Bixler, G. D. & Bhushan, B. Biofouling: lessons from nature. Philos. Trans. R. Soc. Math. Phys. Eng. Sci. 370, 2381–2417 (2012).

6. Chaves Simões, L. & Simões, M. Biofilms in drinking water: problems and solutions. RSC Adv 3, 2520–2533 (2013).

7. Percival, S. L., Suleman, L., Vuotto, C. & Donelli, G. Healthcare-associated infections, medical devices and biofilms: risk, tolerance and control. J. Med. Microbiol. 64, 323–334 (2015).

8. Roberts, A. E. L., Kragh, K. N., Bjarnsholt, T. & Diggle, S. P. The Limitations of In Vitro Experimentation in Understanding Biofilms and Chronic Infection. J. Mol. Biol. 427, 3646–3661 (2015).

9. Carvalho, G., Balestrino, D., Forestier, C. & Mathias, J.-D. How do environment-dependent switching rates between susceptible and persister cells affect the dynamics of biofilms faced with antibiotics? Npj Biofilms Microbiomes 4, (2018).

10. Costerton, J. Introduction to biofilm. Int. J. Antimicrob. Agents 11, 217–221 (1999).

11. Serra, D. O., Richter, A. M., Klauck, G., Mika, F. & Hengge, R. Microanatomy at Cellular Resolution and Spatial Order of Physiological Differentiation in a Bacterial Biofilm. mBio 4, e00103-13–e00103-13 (2013).

12. Ghanbari, A. et al. Inoculation density and nutrient level determine the formation of mushroom-shaped structures in *Pseudomonas aeruginosa* biofilms. Sci. Rep. 6, (2016).

13. Sheraton, M. V. et al. Mesoscopic Energy Minimization Drives *Pseudomonas aeruginosa* Biofilm Morphologies and Consequent Stratification of Antibiotic Activity Based on Cell Metabolism. Antimicrob. Agents Chemother. 62, (2018).

14. Libicki, S. B., Salmon, P. M. & Robertson, C. R. The effective diffusive permeability of a nonreacting solute in microbial cell aggregates. Biotechnol. Bioeng. 32, 68–85 (1988).

15. Hunt, S. M., Werner, E. M., Huang, B., Hamilton, M. A. & Stewart, P. S. Hypothesis for the Role of Nutrient Starvation in Biofilm Detachment. Appl. Environ. Microbiol. 70, 7418–7425 (2004).

16. Stewart, P. S. Diffusion in Biofilms. J. Bacteriol. 185, 1485–1491 (2003).

17. Guélon, T., Mathias, J.-D. & Deffuant, G. Influence of spatial structure on effective nutrient diffusion in bacterial biofilms. J. Biol. Phys. 38, 573–588 (2012).

18. Rudge, T. J., Steiner, P. J., Phillips, A. & Haseloff, J. Computational Modeling of Synthetic Microbial Biofilms. ACS Synth. Biol. 1, 345–352 (2012).

19. Rudge, T. J., Federici, F., Steiner, P. J., Kan, A. & Haseloff, J. Cell Polarity-Driven Instability Generates Self-Organized, Fractal Patterning of Cell Layers. ACS Synth. Biol. 2, 705–714 (2013).

20. Blanchard, A. E. & Lu, T. Bacterial social interactions drive the emergence of differential spatial colony structures. BMC Syst. Biol. 9, (2015).

21. Smith, W. P. J. et al. Cell morphology drives spatial patterning in microbial communities. Proc. Natl. Acad. Sci. 114, E280–E286 (2017).

22. Goldschmidt, F., Regoes, R. R. & Johnson, D. R. Successive range expansion promotes diversity and accelerates evolution in spatially structured microbial populations. ISME J. 11, 2112 (2017).

23. Jauffred, L., Vejborg, R. M., Korolev, K. S., Brown, S. & Oddershede, L. B. Chirality in microbial biofilms is mediated by close interactions between the cell surface and the substratum. ISME J. 11, 1688 (2017).

24. Eriksen, R. S., Svenningsen, S. L., Sneppen, K. & Mitarai, N. A growing microcolony can survive and support persistent propagation of virulent phages. Proc. Natl. Acad. Sci. 115, 337–342 (2018).

25. Xiao, J. et al. Biofilm three-dimensional architecture influences in situ pH distribution pattern on the human enamel surface. Int. J. Oral Sci. 9, 74–79.

26. Liu, J. et al. Coupling between distant biofilms and emergence of nutrient time-sharing. Science 356, 638–642 (2017).

27. McConnell, G. et al. A novel optical microscope for imaging large embryos and tissue volumes with sub-cellular resolution throughout. eLife 5, e18659 (2016).

28. McConnell, G. & Amos, W. B. Application of the Mesolens for subcellular resolution imaging of intact larval and whole adult *Drosophila*. J. Microsc. 270, 252–258 (2018).

29. Schniete, J. et al. Fast Optical Sectioning for Widefield Fluorescence Mesoscopy with the Mesolens based on HiLo Microscopy. Sci. Rep. 8, (2018).

30. Drescher, K. et al. Architectural transitions in *Vibrio cholerae* biofilms at single-cell resolution. Proc. Natl. Acad. Sci. 113, E2066–E2072 (2016).

31. Yan, J., Sharo, A. G., Stone, H. A., Wingreen, N. S. & Bassler, B. L. Vibrio cholerae biofilm growth program and architecture revealed by single-cell live imaging. Proc. Natl. Acad. Sci. 113, E5337–E5343 (2016).

32. Hartmann, R. et al. Emergence of three-dimensional order and structure in growing biofilms. Nat. Phys. 15, 251–256 (2019).

33. Lagree, K., Desai, J. V., Finkel, J. S. & Lanni, F. Microscopy of fungal biofilms. Curr. Opin. Microbiol. 43, 100–107 (2018).

34. Xiao, J. et al. Biofilm three-dimensional architecture influences in situ pH distribution pattern on the human enamel surface. Int. J. Oral Sci. 9, 74–79 (2017).

35. Thomsen, H. et al. Delivery of cyclodextrin polymers to bacterial biofilms — An exploratory study using rhodamine labelled cyclodextrins and multiphoton microscopy. Int. J. Pharm. 531, 650–657 (2017).

36. Shemesh, H. et al. High frequency ultrasound imaging of a single-species biofilm. J. Dent. 35, 673–678 (2007).

37. Vaidya, K., Osgood, R., Ren, D., Pichichero, M. E. & Helguera, M. Ultrasound Imaging and Characterization of Biofilms Based on Wavelet De-noised Radiofrequency Data. Ultrasound Med. Biol. 40, 583–595 (2014).

38. Xi, C., Marks, D., Schlachter, S., Luo, W. & Boppart, S. A. High-resolution three-dimensional imaging of biofilm development using optical coherence tomography. J. Biomed. Opt. 11, 034001 (2006).

39. Wagner, M., Taherzadeh, D., Haisch, C. & Horn, H. Investigation of the mesoscale structure and volumetric features of biofilms using optical coherence tomography. Biotechnol. Bioeng. 107, 844–853 (2010).

40. Leite de Andrade, M. C. et al. A new approach by optical coherence tomography for elucidating biofilm formation by emergent Candida species. PLOS ONE 12, e0188020 (2017).

41. Drury, W. J., Characklis, W. G. & Stewart, P. S. Interactions of 1 μm latex particles with *Pseudomonas aeruginosa* biofilms. Water Res. 27, 1119–1126 (1993).

42. Stoodley, P., Lewandowski, Z. & others. Liquid flow in biofilm systems. Appl. Environ. Microbiol. 60, 2711–2716 (1994).

43. Asally, M. et al. Localized cell death focuses mechanical forces during 3D patterning in a biofilm. Proc. Natl. Acad. Sci. 109, 18891–18896 (2012).

44. Wilking, J. N. et al. Liquid transport facilitated by channels in Bacillus subtilis biofilms. Proc. Natl. Acad. Sci. 110, 848–852 (2013).

45. Kempes, C. P., Okegbe, C., Mears-Clarke, Z., Follows, M. J. & Dietrich, L. E. P. Morphological optimization for access to dual oxidants in biofilms. Proc. Natl. Acad. Sci. 111, 208–213 (2014).

46. Jo, J., Cortez, K. L., Cornell, W. C., Price-Whelan, A. & Dietrich, L. E. An orphan cbb3-type cytochrome oxidase subunit supports Pseudomonas aeruginosa biofilm growth and virulence. 30 (2017).

47. Nuñez, I. N. et al. Artificial Symmetry-Breaking for Morphogenetic Engineering Bacterial Colonies. ACS Synth. Biol. 6, 256–265 (2017).

48. Wimpenny, J. W. T. & Coombs, J. P. Penetration of oxygen into bacterial colonies. Microbiology 129, 1239–1242 (1983).

49. Peters, A. C., Wimpenny, J. W. T. & Coombs, J. P. Oxygen Profiles in, and in the Agar Beneath, Colonies of *Bacillus cereus*, *Staphylococcus albus* and *Escherichia coli*. J. Gen. Microbiol. 133, 1257–1263 (1987).

50. Jeanson, S., Floury, J., Gagnaire, V., Lortal, S. & Thierry, A. Bacterial Colonies in Solid Media and Foods: A Review on Their Growth and Interactions with the Micro-Environment. Front. Microbiol. 6, (2015).

51. Hwang, G. et al. Simultaneous spatiotemporal mapping of in situ pH and bacterial activity within an intact 3D microcolony structure. Sci. Rep. 6, (2016).

52. Webb, J. S. et al. Cell Death in *Pseudomonas aeruginosa* Biofilm Development. J. Bacteriol. 185, 4585–4592 (2003).

53. Moscona, A. A. Aggregation of sponge cells: Cell-linking macromolecules and their role in the formation of multicellular systems. In Vitro 3, 13–21 (1967).

54. Lavrov, A. I. & Kosevich, I. A. Sponge cell reaggregation: Mechanisms and dynamics of the process. Russ. J. Dev. Biol. 45, 205–223 (2014).

55. Xu, H., Dauparas, J., Das, D., Lauga, E. & Wu, Y. Self-organization of swimmers drives long-range fluid transport in bacterial colonies. Nat. Commun. 10, (2019).

56. Jolivet-Gougeon, A. & Bonnaure-Mallet, M. Biofilms as a mechanism of bacterial resistance. Drug Discov. Today Technol. 11, 49–56 (2014).

57. Elbing, K. L. & Brent, R. Recipes and Tools for Culture of Escherichia coli. Curr. Protoc. Mol. Biol. 125, e83 (2019).

58. Dempster, J., Wokosin, D. L., McCloskey, K. D., Girkin, J. M. & Gurney, A. M. WinFluor: an integrated system for the simultaneous recording of cell fluorescence images and electrophysiological signals on a single computer system. Br. J. Pharmacol. 137, 146 (2002).

59. Lambertsen, L., Sternberg, C. & Molin, S. Mini-Tn7 transposons for site-specific tagging of bacteria with fluorescent proteins. Environ. Microbiol. 6, 726–732 (2004).

60. Schindelin, J. et al. Fiji: an open-source platform for biological-image analysis. Nat. Methods 9, 676–682 (2012).

