## Supplementary Information for "Functional channels in mature *E. coli* colonies"

### A network of intra-colony channels facilitates nutrient acquisition in mature *Escherichia coli* biofilms

**This PDF file includes:**

Supplementary figures S1 to S3
Table S1

### Supplementary Information

#### Supplementary Figures

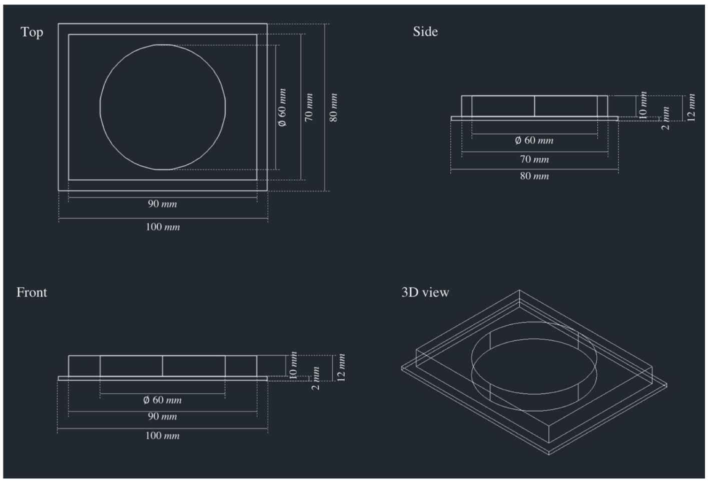


Supplementary Figure 1. **Schematic for bacterial culture and imaging mould.** Technical drawing shows dimensions for the 3D-printed imaging mould used in this study. The design was based on a Petri dish-come-microscope slide with a central well where bacteria can be cultured as required. The top surface could be sealed with a large glass coverslip which was custom-made for the Mesolens, and if required, the design allows the imaging chamber to be incorporated into an immersion bath.

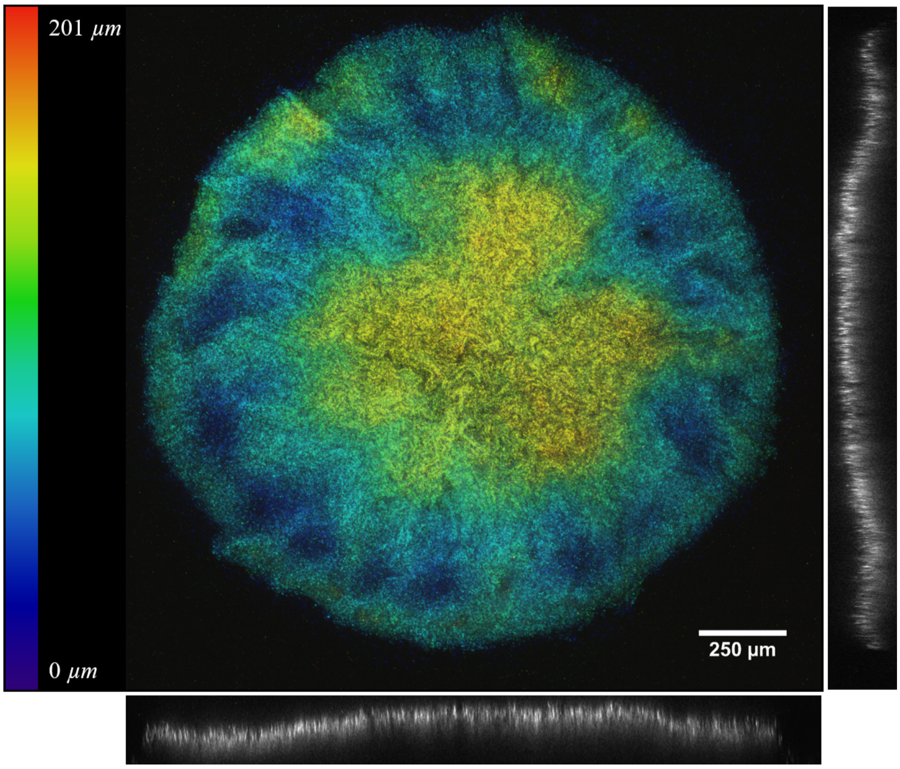


Supplementary Figure 2. **Confocal laser scanning mesoscopy revealed that intra-colony spatial patterns are not an artefact of the deconvolution process.** A maximum intensity projection of a single JM105-miniTn7-*gfp* colony measuring 201 μm thick is shown with a colour coded LUT applied to each optical section depending on its axial position within the dataset. Intra-colony channels could be observed throughout the colony and converged towards the apex of the biofilm. Accompanying orthogonal sections are presented from a cross-section at the centre of the colony. Channels were observed to pass through the majority of the colony from the base to the apex suggesting that they originate at the base of the biofilm.

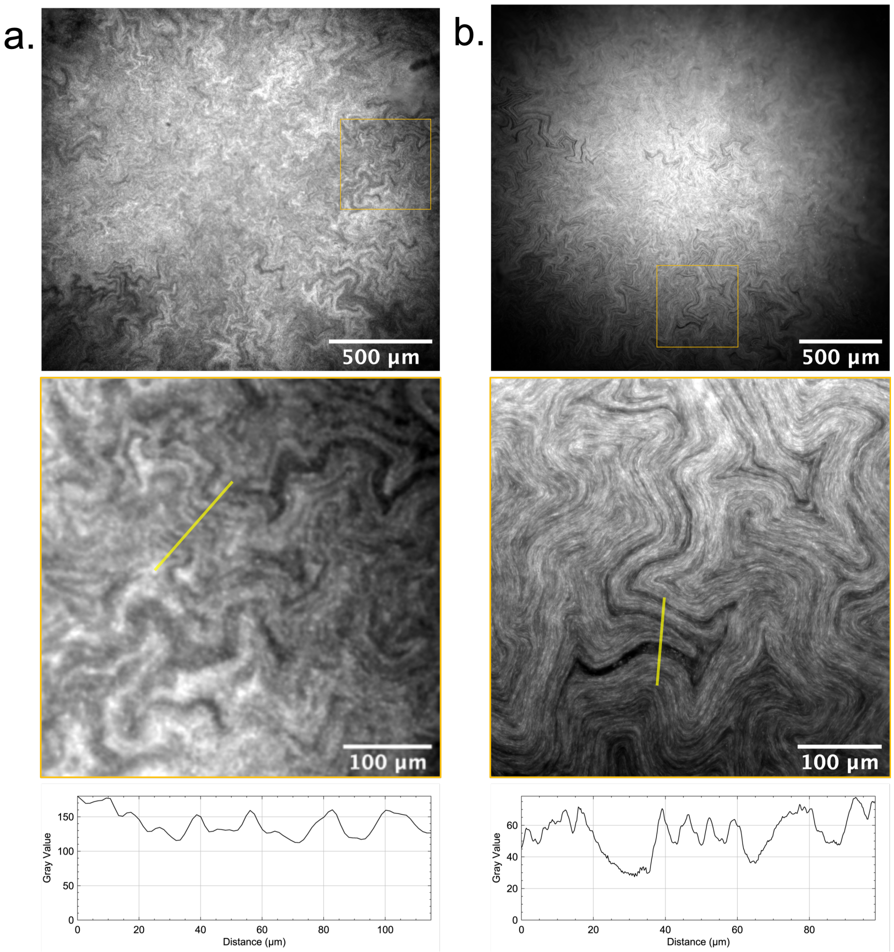


Supplementary Figure 3. **Comparison of conventional low-NA widefield epi-fluorescence microscopy and high-NA mesoscopy. (a)** Image of a JM105-miniTn7-*gfp* macro-colony biofilm acquired using a conventional low-magnification, low-NA objective lens on an upright widefield epi-fluorescence microscope. **(b)** A mature JM105-miniTn7-*gfp* colony biofilm imaged using widefield epi-fluorescence mesoscopy. Intracolony spatial patterns are evident in both (a) and (b). Magnified regions have comparable FOVs, and line ROIs, shown in yellow, indicate that the increased NA of the Mesolens results in spatial higher resolution than a low-magnification, low numerical aperture lens.

Supplementary Table 1. **List of bacterial strains**

| Strain | Characteristics | Source |
|---|---|---|
| JM105 | Routine K12-derived laboratory strain | DSMZ, Germany |
| JM105-miniTn7-*gfp* | miniTn7-*gfp* | [70] |
| JM105-miniTn7-*HcRed1* | miniTn7-*HcRed1* | [70] |
| JM105-pJM058 | $P_{BAD}$-*gfp* | Gifted from Nicola Holden (James Hutton Institute, UK) |

Supplementary Movie 1. **3D reconstruction of an *E. coli* macro-colony biofilm using confocal mesoscopy.**
